# Single-cell analysis reveals distinct immune characteristics of hepatocellular carcinoma in HBV-positive versus HBV-negative cases

**DOI:** 10.1101/2024.12.30.630656

**Authors:** Ke Liu, Erbao Chen, Jiaming Liang, Yanyan Li, Binghua Cheng, Wenli Shi, Zeyu Zhou, Wenjie Zhou, Hui Tian, Dongye Yang, Ximing Shao, Hongchang Li

## Abstract

Infection with the Hepatitis B virus (HBV) is a key risk factor for Hepatocellular carcinoma (HCC) development and progression. It is widely recognized that immunopathological mechanisms are pivotal in developing HBV-related HCC. Nevertheless, the specific mechanisms by which HBV-induced modifications within the tumor microenvironment (TME) contribute to HCC pathogenesis are still not well understood. Here, we utilized single-cell RNA sequencing to analyze and compare the immune landscapes between HBV-positive and HBV-negative HCC. We discovered that HBV infection significantly modifies the immune cell makeup and state, and leads to the suppression and exhaustion of T cells within the TME. Specifically, an increase in SLC4A10+ CD8+ T cells and IFITM3+ macrophages was observed, along with the upregulation of the gene SLC35F1 in various immune cell subtypes. These findings offer valuable insights into the alteration of the immunological microenvironment in HCC associated with HBV infection, suggesting possible targets for immunotherapeutic intervention.

## Introduction

Hepatocellular carcinoma (HCC), a primary liver cancer, is a significant global health issue, ranking as the sixth most frequent cancer and the third leading cause of cancer-related mortality around the world (1, 2). The incidence of HCC is particularly high in East Asia and sub-Saharan Africa, regions where chronic hepatitis B virus (HBV) infection is endemic(3). HBV is recognized as a significant risk factor in the emergence of HCC. Approximately 300 million individuals are affected by HBV globally, with the virus accounting for nearly 50% of all HCC cases and virtually all childhood HCC cases(4). Understanding the mechanisms by which HBV infection induces HCC is critical for developing effective prevention and treatment strategies, particularly for high-risk populations.

One of the critical aspects of HBV infection is its profound impact on the immune microenvironment. The virus has evolved various strategies to evade the host immune system, enabling it to maintain a chronic infection. This evasion is partly achieved by altering the immune microenvironment, which includes the recruitment and modulation of immune cells like natural killer (NK) cells, T cells, and macrophages. These changes can lead to an impaired immune response, enabling the virus to persist and cause ongoing liver damage(5, 6).

The immune microenvironment is vital in cancer development. In the context of HBV infection, the altered immune landscape can contribute to the onset and progression of liver cancer. For instance, there is often a rise in myeloid-derived suppressor cells (MDSCs) and regulatory T cells (Tregs), which hinder effective immune responses and support immune tolerance(7, 8). Conversely, the activity of natural killer (NK) cells and cytotoxic T lymphocytes (CTLs), which are crucial for clearing viral infections, is often diminished(9, 10). This dysregulation of immune cell populations can create a tumor-promoting microenvironment. Despite significant advancements in medical research(11-14), our understanding of the immune mechanisms by which HBV infection promotes HCC remains limited, largely because of the intricate nature of the immune environment.

Single-cell RNA sequencing (scRNA-seq) has brought about a revolutionary change in how the immune system is comprehensively analyzed. Its application to immune cell populations has enabled an in-depth characterization of the immune microenvironment and the identification of novel immune subsets in various tumor types, such as lung, breast, colorectal, liver, and pancreatic cancers(15-20). In this study, we collected tumors from HCC patients who were either HBV-positive or HBV-negative, conducted scRNA-seq, and compared the immune microenvironment of HBV-positive and HBV-negative tumors. We found that HBV infection significantly alters the landscape of immune cells within HCC, and leads to a pronounced suppression and exhaustion of T cell. Specifically, we observed an increased proportion of SLC4A10+ CD8+ T cells, which possess the capacity to facilitate tumor initiation, growth, and metastasis, alongside a heightened presence of IFITM3+ macrophages, which play critical roles in viral clearance. Additionally, we identified a gene, SLC35F1, that is significantly upregulated across various immune cell subtypes within HBV-infected tumor. These findings deliver valuable immunological insights into the pathogenesis and progression of liver cancer in the setting of HBV infection and offer potential targets for immunotherapy in HCC.

## Materials and methods

### Patient specimens

Enrolled in this study were six patients from Peking University Shenzhen Hospital who underwent liver resection and received a pathological diagnosis of HCC between 2022 and 2024. All patients had not received any anti-tumor therapies before their surgeries. From each individual, fresh tumor samples and their corresponding para-cancerous tissues were gathered.

### scRNA-seq library preparation

scRNA-seq libraries were generated using the SeekOne® Digital Droplet Single Cell 3’ Library Preparation Kit (Cat# K00202, SeekGene). The libraries were then sequenced on an Illumina NovaSeq 6000 platform with a paired-end 150 bp (PE150) read length.

### Raw data processing

Data preprocessing was conducted using the Seeksoultools suite with its standard settings. Initially, the process involved converting binary base call files into FASTQ format through the use of cellranger mkfastq, which referenced a sample sheet that incorporated Seekgene barcodes. Subsequently, the seeksoultools count pipeline facilitated the alignment of sequence reads to the reference genome supplied by 10x Genomics (specifically, the human reference dataset refdata-gex-GRCh38-2020-A). This alignment process included the tallying of reads for each gene, with intronic reads being factored into the count matrix. Additionally, it encompassed the computation of clustering and summary statistics. The output from the seeksoulgene count for each sample was pooled, ensuring uniformity in sequencing depth across samples. Finally, the feature-barcode matrices were recalculated using the seeksoultools package version 1.2.2.

### Single-cell gene expression quantification

To accurately measure gene expression, we implemented a strategy that only considered unique molecular identifiers (UMIs) once, thereby preventing the overcounting of PCR-amplified transcripts. This approach led to the creation of cell-gene UMI matrices, which were crucial for further analysis. The process included the elimination of cells with extreme UMI counts—those with very high (over 6000) or very low (under 200) counts—to minimize the impact of unwanted variability and to enhance the quality of the cell data. Additionally, to mitigate the influence of potential doublets, we employed a doublet detection method to identify and exclude such cells from our dataset.

### Quality control and batch correction

In order to enhance the quality of our cell analysis and eliminate both low-quality cells and doublets, we applied specific criteria for each sample. Cells were excluded if they had less than 200 unique molecular identifiers (UMIs) or if they expressed fewer than 200 or more than 8000 genes. This step was crucial for removing cells that might not be reliable for analysis. Furthermore, to exclude cells that might be dead or in the process of dying, we removed those with over 5% of their UMIs derived from the mitochondrial genome. After these filtrations, a total of 121,848 high-quality single-cell transcriptomes were obtained from all samples.For the integration of samples from different tissues or patients, we utilized the harmony function from the R package Harmony (version 1.2.1) to perform batch correction. This step was essential for harmonizing the data from various sources. The counts that were corrected using harmony were subsequently employed for Unsupervised Uniform Manifold Approximation and Projection (UMAP) analysis in Seurat, which is a powerful tool for dimensionality reduction and visualization of complex datasets.

### Cells clustering

For the purpose of cell clustering, we initially built a K-nearest neighbor (KNN) graph from a Euclidean distance matrix within the principal component analysis (PCA) space, which was then transformed into a shared nearest neighbor (SNN) graph to identify tightly connected cell communities. The Louvain algorithm was employed to cluster cells, aiming to optimize modularity. For data visualization, we applied the Unsupervised Uniform Manifold Approximation and Projection (UMAP) technique to the cell loadings of selected principal components (PCs), incorporating the cluster assignments derived from the graph-based clustering process. When dealing with more than two clusters, we utilized the “find_all_markers” function in Seurat, setting the log fold change threshold to 0.5 and employing the Wilcoxon rank-sum test to pinpoint marker genes specific to each cluster. For the purpose of identifying genes that are differentially expressed between any two clusters, we employed Seurat’s “find.markers” function, also with a log fold change threshold of 0.5 and using the Wilcoxon rank-sum test. The Seurat R package version 5.1.0 was employed to conduct all the analyses detailed in this section.

### Gene sets enrichment analysis

Functional enrichment analysis of differentially expressed genes (DEGs) at P < 0.05 was executed utilizing the R package clusterProfiler version 4.12.6. Additionally, Gene Set Enrichment Analysis (GSEA) was executed with the aid of GSEA desktop application, sourcing gene sets and molecular signatures from the Molecular Signatures Database. The analysis involved 1000 permutations of gene sets to derive normalized enrichment scores, with a significance threshold of P-value < 0.05 applied to identify enriched results.

### Transcription factor regulon analysis

The regulatory network and regulon activity were analyzed using pySCENIC. Regulon activity, quantified by AUC, was assessed through the AUCell module in pySCENIC, with active regulons identified based on the module’s default threshold. Differentially expressed regulons were determined using the Wilcoxon rank-sum test, implemented via the “FindAllMarkers” function of the Seurat R package, with designated parameters: logfc.threshold = 0.25, min.pct = 0.1, pseudocount.use = F, only.pos = T. A heatmap was generated to visualize the scaled regulon activity expression.

### TCGA-LIHC survival analysis

For TCGA LIHC data, gene expression and clinical information were obtained from The Cancer Genome Atlas (TCGA) through GDC portal (https://portal.gdc.cancer.gov). Gene expression TPM values were log-transformed for analysis. Survival analysis was performed with the R package *survival*, while the *surv_cutpoint()* function from the *survminer* package was employed to identify optimal cutpoints for cell infiltration, categorizing the data into two groups. Kaplan–Meier survival curves were ploted with the *survfit()* function, and statistical significance among groups was determined by employing the log-rank test.

### Cell–cell communication analysis

Intercellular communication in HBV-infected versus non-HBV-infected HCC was analyzed using the CellChat R package. Separate CellChat objects were created for each condition and merged via the *mergeCellChat* function. Interaction counts and strengths were assessed with *compareInteractions*, while differences in interaction numbers and strengths between cell populations across the two datasets were visualized in a heatmap using *netVisual_heatmap*. Signaling pathway distances were calculated with *rankSimilarity*, and significant changes in signal sending and receiving for each cell type were identified using *netAnalysis_signalingRole_scatter*. Specific cell population signaling changes were further examined with *netAnalysis_signalingChanges_scatter*. Finally, communication dynamics for individual signaling pathways were compared using the *netVisual_aggregate* function.

## Results

### scRNA-seq and cell typing in primary HCC and paired non-tumor liver tissues

To construct a single-cell atlas of HCC, we recruited 6 patients with primary tumors. The patients, aged 49 to 75 and including both males and females, exhibited no vascular invasion or metastatic features and were classified as TNM stage I (Fig. S1A). We profiled 121,848 single cells, with an average detection of 1,892 genes per cell (Fig. S1B). To map the global cellular microenvironment of HCCs, scRNA-seq data from all tissues and patients were integrated through a batch correction approach based on the *harmony* package. Unsupervised clustering with a shared-nearest neighbor (SNN) method identified 11 distinct cell clusters (Fig. S1C). These clusters were annotated using canonical marker genes, revealing hepatocyte, T/NK cell, B cell, myeloid, endothelial, and fibroblast clusters (Fig. 1A, B, and Fig. S1D). Marker gene expression further validated the specificity of each cluster (Fig. 1C). We also analyzed the distribution of annotated cell types across the 12 samples (Fig. 1D). Notably, the percentage of T/NK cells was considerably lower in tumor tissues than in adjacent tissues, suggesting an immune-suppressive tumor microenvironment (Fig. 1E).

**Fig. 1.**
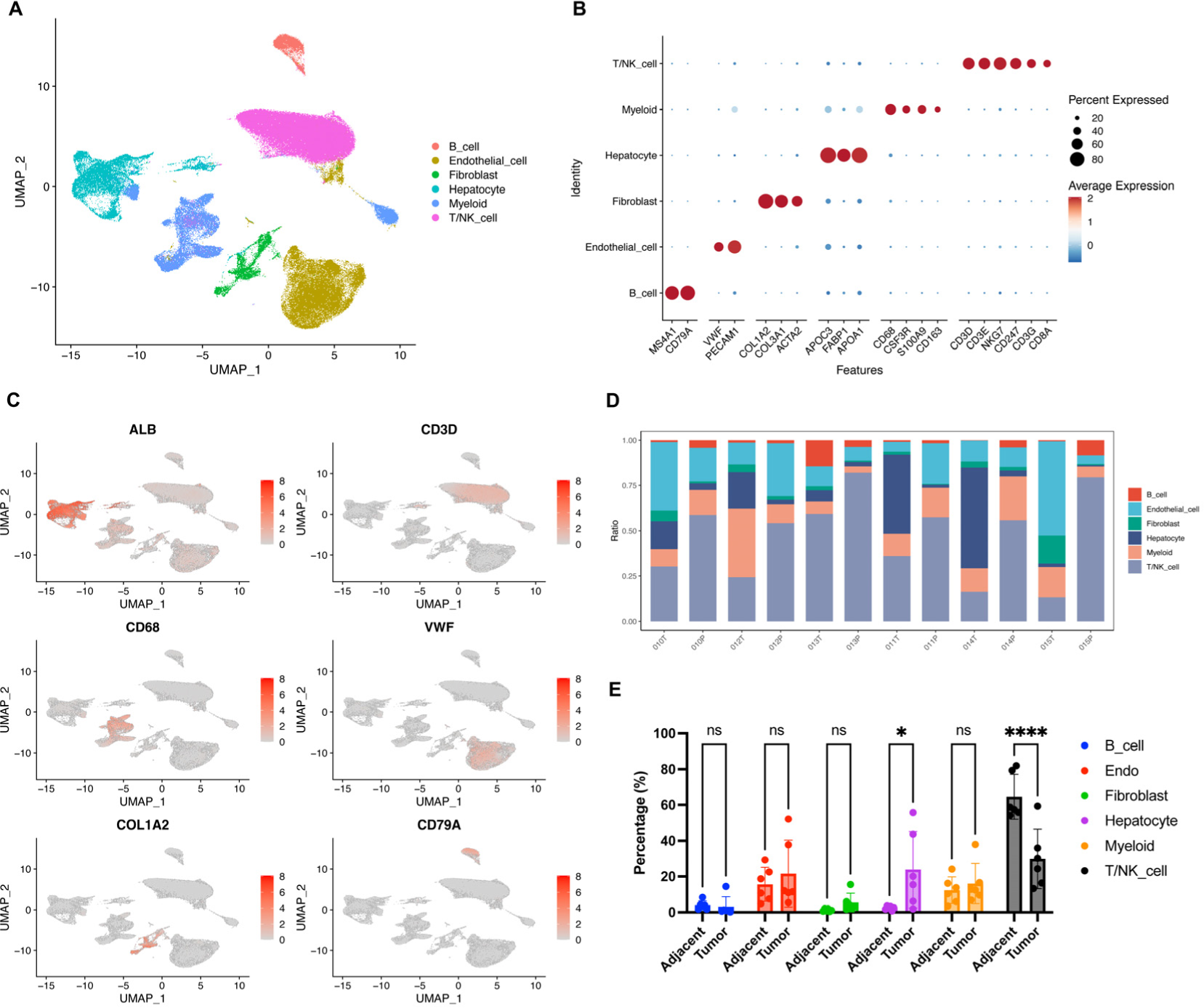
scRNA-seq analysis of primary HCC and adjacent non-tumor liver tissues. (A) UMAP plot displaying the distribution of cells from HCC and adjacent tissues in 6 patients. (B) Dot plot illustrating the percentage of cells expressing canonical marker genes alongside their average expression levels across major cell types. (C) UMAP plots showing the expression patterns of signature genes for six major cell types, with colors indicating expression levels. (D) Stacked bar plots representing the proportions of major cell types in each sample, arranged by tissue type (T: tumor; P: para-tumor). (E) Quantification of cell type percentages in tumor and adjacent tissues.

### HBV-infected HCC displays unique clustering and transcriptomic profile compared with non-HBV-infected HCC

Besides isolated tumor and adjacent tissues, our scRNAseq dataset also includes HBV-infection or non-HBV-infection information, which may provide us with useful resources to compare the clustering and transcriptomic profile of HCC with or without HBV-infection. To delve into this issue, we focus on our tumor-scRNAseq dataset and characterize the cell clustering information. To notice, here we defined the HBV-infected HCC as HBV-positive or Po, and defined the non-HBV-infected HCC as HBV-negative or Ne. The cell-density-map showed that there is clear difference between the HBV-negative and HBV-positive samples (Fig.2A). Most evidently, the proportions of B cells and T/NK cells are dramatically increased under HBV-infected condition, which suggest HBV-infection may result in the increased infiltration or proliferation of those cells (Fig.2A). To further confirm this observation, we summarized the percentage of each cell type of 6 samples. Although, there seems variations among HBV-negative or HBV-positive samples, possibly due to the heterogeneity of patients age, status or other conditions, the overall trend of increased percentage of B cells and T/NK cells could be steadily observed (Fig.2B). Moreover, the calculation of average percentage of cell types also showed B cells and T/NK cells increased in HBV-infected tumors (Fig.2C). Collectively, we conclude that the HBV-infected HCC may harbor increased T/NK cells and B cells when making comparison with non-HBV-infected tumors.

**Fig. 2.**
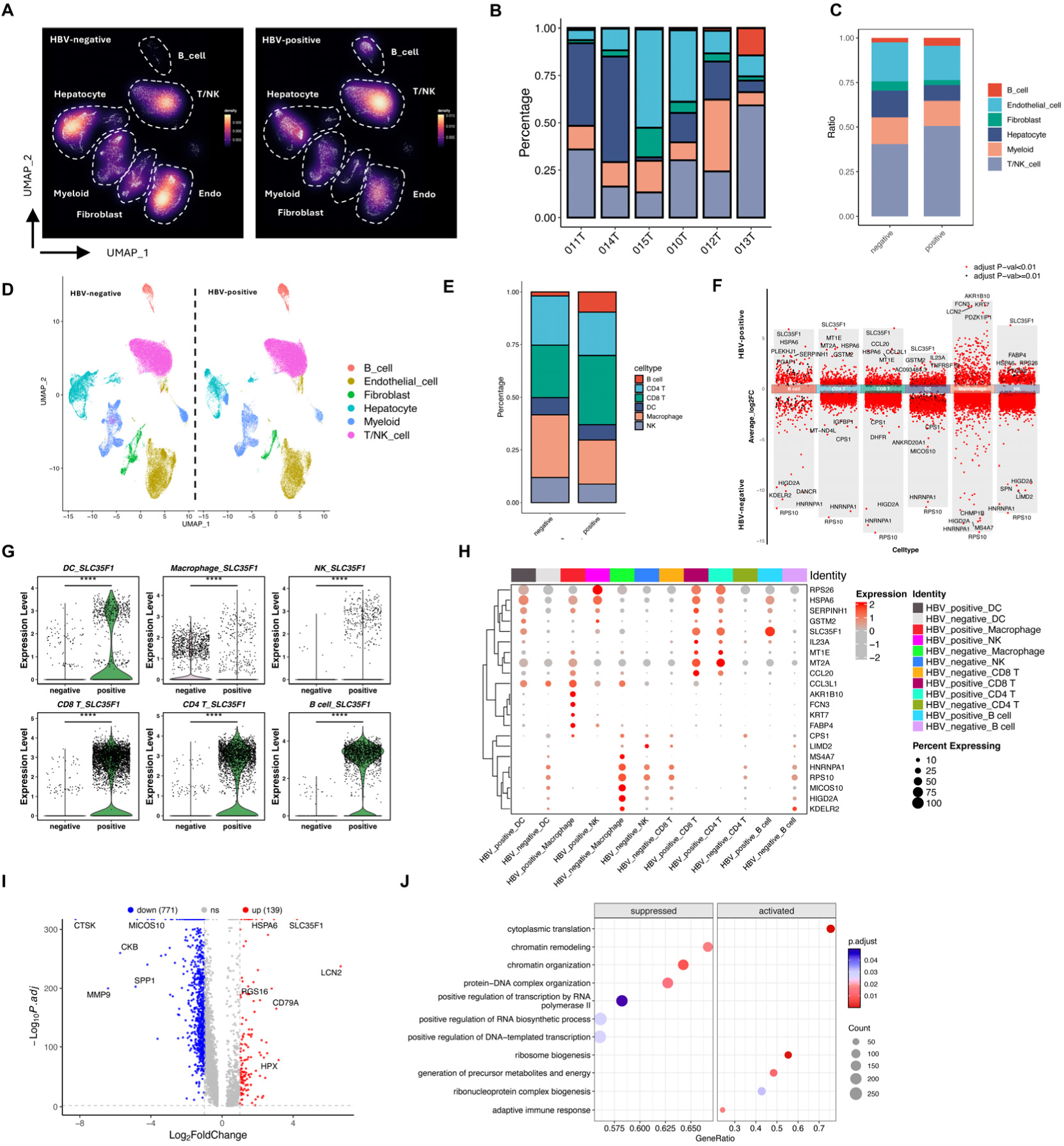
scRNA-seq profiling of HBV-infected and non-HBV-infected HCC. (A) Density UMAP plots illustrating the cell density of clusters in HBV-negative (non-HBV-infected) and HBV-positive (HBV-infected) samples.(B) Stacked bar plots displaying the percentages of major cell types in HBV-negative and HBV-positive samples. (C) Summarized stacked bar plots showing the overall percentages of major cell types in HBV-negative and HBV-positive samples. (D) UMAP plots depicting further clustering of immune cells in HBV-negative and HBV-positive samples, respectively. (E) Quantification of the percentage of selected major immune cell types in HBV-negative and HBV-positive samples. (F) Volcano plot showing differentially expressed genes (DEGs) by comparing immune cells from HBV-negative and HBV-positive samples. (G) Violin plots illustrating *SLC35F1* expression across immune cell types with or without HBV infection. (H) Dot plot showing the top 5 DEGs for each immune cell type under HBV-negative and HBV-positive conditions. (I) Volcano plot displaying overall DEGs by combining all immune cells for a comparative analysis. (J) GSEA plot indicating activated and suppressed Gene Ontology (GO) terms in immune cells with or without HBV infection.

Myeloid cells include various subtypes such as macrophages, neutrophils, and dendritic cells (DCs). To explore the dynamics of specific immune cells in response to HBV infection, we conducted further clustering on the identified cell types. Through marker gene expression analysis, we identified CD4 T cells, CD8 T cells, dendritic cells, macrophages, monocytes, neutrophils, and NKT cells (Fig. 2D and Fig. S2A, B). Among major immune cells, B cells, CD4 T cells, CD8 T cells, DCs, macrophages, and NK cells exhibited significant proportional differences between HBV-negative and HBV-positive samples (Fig. 2E). Specifically, HBV infection led to an overall increase in the proportion of B cells and CD8 T cells, while the percentages of NK cells and macrophages decreased. These findings suggest diverse responses of immune cell types to HBV infection. To investigate molecular changes within these immune cells under HBV infection, we conducted differential gene expression (DEG) analysis. This revealed both upregulated and downregulated genes in each type of immune cell (Fig. 2F). Notably, SLC35F1, a gene associated with nucleoside transport and metabolism(21, 22), was consistently upregulated across multiple immune cells, including B cells, CD4 T cells, CD8 T cells, DCs, and NK cells (Fig. 2F). Further examination confirmed the significant upregulation of SLC35F1 in all immune cell types, including macrophages (Fig. 2G). When comparing the top four DEGs for each immune cell type, we observed both distinct and shared DEGs, with SLC35F1 prominently highlighted (Fig. 2H). Additionally, we also combined all the immune cells in HBV-negative or HBV-positive conditions respectively and performed DEG analysis. This revealed 139 upregulated and 771 downregulated genes upon the HBV-infection (Fig. 2I). Gene ontology (GO) and Kyoto Encyclopedia of Genes and Genomes (KEGG) analyses of these altered genes showed that upregulated genes were enriched in pathways related to ribosome biogenesis, protein folding, and leukocyte activation, suggesting that HBV infection promotes active protein synthesis, metabolism, and immune activation (Fig. S2C-D). Conversely, downregulated genes were associated with insulin response, hormone signaling, and AMPK activation pathways (Fig. S2C-D). Gene set enrichment analysis (GSEA) further revealed that immune cells in HBV-infected tumors exhibited increased signaling pathways related to adaptive immune responses and antiviral defense (Fig. 2J and Fig. S2E). Together, our scRNA-seq analysis demonstrates that HBV infection alters both the proportion and transcriptomic profiles of immune cells, with SLC35F1 identified as a commonly upregulated gene across most immune cell types in response to infection.

### Characterization of T Cell heterogeneity in both HBV-infected and non-HBV-infected HCC

T cell immunity plays a vital role in both tumor growth and immune surveillance in HCC(23, 24). Infection with HBV is known to significantly reshape the immune landscape, impacting the effectiveness of immune responses(6, 25). However, the heterogeneity of T cells of HCC under different HBV infection status is still poorly investigated. To understand how HBV infection influences T cell diversity in HCC, we analyzed our scRNA-seq data with a focus on CD4+ and CD8+ T cells.

CD4+ T cells is crucial for orchestrating immune responses. Leveraging established cellular signature markers from the literature, we classified CD4+ T cells into six distinct subtypes to examine their proportional differences between HBV-infected and non-HBV-infected samples (Fig. 3A, B, and Fig. S3A). These subtypes were further defined by their marker gene expression profiles to elucidate their functional roles (Fig. 3C). Our analysis revealed a decrease in the proportion of IL7R+ CD4+ T cells and an increase in ANXA1+ CD4+ T cells in HBV-infected samples (Fig. 3B). This suggests a potential link between these T cell subtypes and HBV infection status. To investigate further, we conducted a DEG analysis for each CD4+ T cell subtype between HBV-positive and HBV-negative groups. Clearly, SLC35F1 was consistently upregulated across all classified subtypes (Fig. 3D), highlighting its potential role as a key mediator or functional gene responsive to HBV infection.

**Fig. 3.**
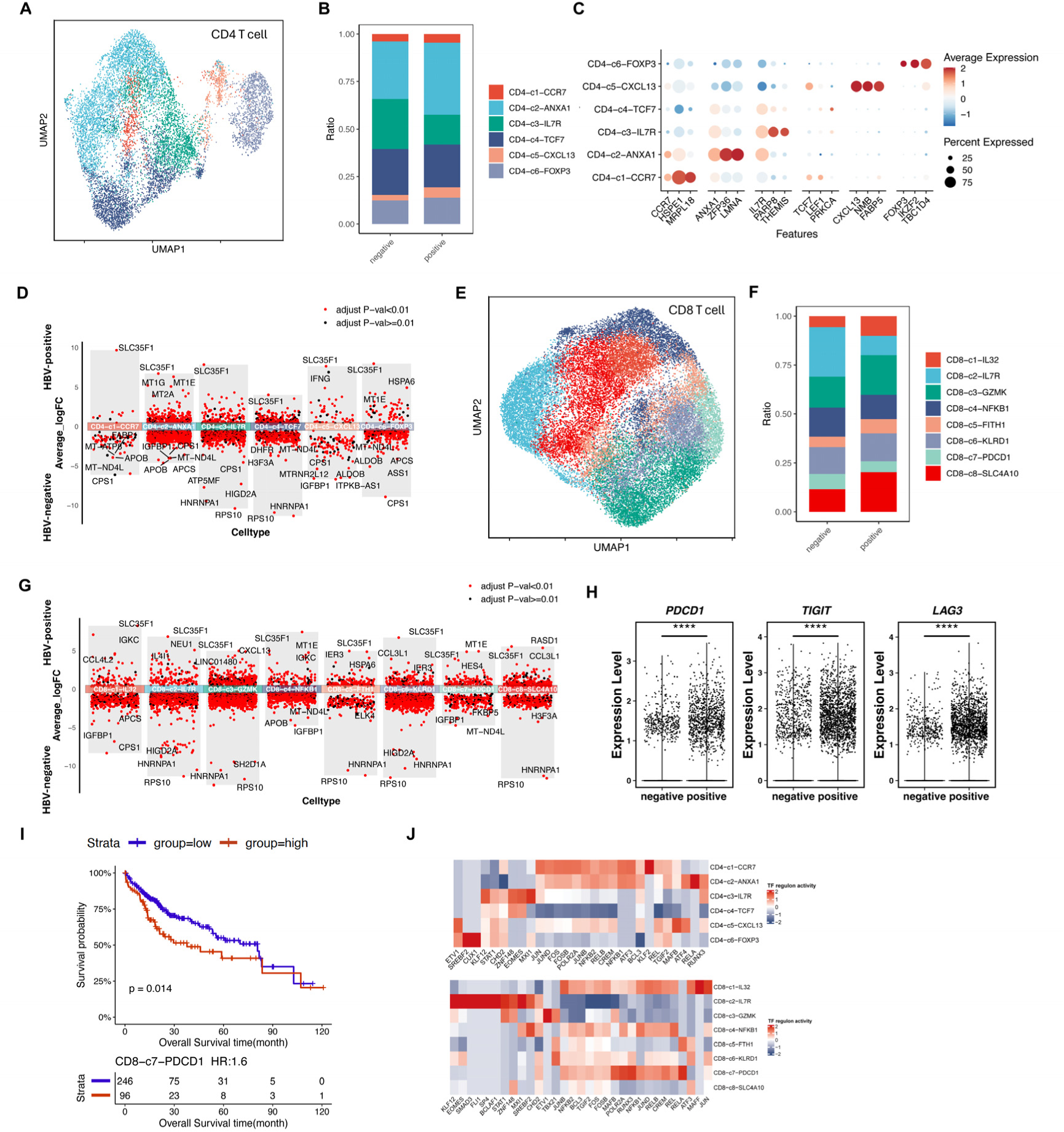
Analysis of T cell heterogeneity. (A) UMAP plot of CD4+ T cells, with cell subtypes distinguished by color. (B) Stacked bar plots summarizing the proportions of CD4+ T cell subtypes in HBV-negative and HBV-positive samples. (C) Dot plot showing the percentages of cells expressing marker genes alongside their average expression levels across CD4+ T cell subtypes. (D) Volcano plot highlighting DEGs between HBV-negative and HBV-positive CD4+ T cells. (E) UMAP plot of CD8+ T cells, with cell subtypes color-coded. (F) Stacked bar plots summarizing the proportions of CD8+ T cell subtypes in HBV-negative and HBV-positive samples. (G) Volcano plot of DEGs comparing HBV-negative and HBV-positive CD8+ T cells. (H) Violin plot depicting the expression of exhaustion marker genes in HBV-negative and HBV-positive CD8+ T cells. (I) Kaplan–Meier curve illustrating survival of LIHC patients with varying levels of CD8-c7-PDCD1 infiltration. (J) Heatmap showing normalized TF regulon activity in CD4+ and CD8+ T cell subtypes as predicted by pySCENIC.

Besides CD4 T cells, we also took care about the CD8+ T cells, which is crucial for cytotoxic responses against tumors. We classified CD8+ T cells into eight subtypes (Fig.3E) and observed distinct differences in subtype distributions between HBV-infected and non-infected groups (Fig.3F and Fig.S3B). To understand their functional states, we assessed key marker genes indicative of exhaustion, effector, naive, and memory characteristics (Fig.S3C). The subtypes of CD8+ T cells exhibited more dynamic changes in response to HBV infection (Fig. 3F). Notably, the proportion of SLC4A10+ CD8+ T cells increased, while that of PDCD1+ CD8+ T cells decreased under HBV-infected conditions. Furthermore, we performed DEG analysis to examine the differences in the transcriptomic profiles of each CD8+ T cell subtype between HBV-negative and HBV-positive groups. Again, SLC35F1 emerged as a prominently upregulated gene across all subtypes, indicating its potential regulatory role in CD8+ T cells (Fig. 3G).

Interestingly, despite the reduced proportion of PDCD1+ CD8+ T cells under HBV infection, we found a significant upregulation of exhaustion marker genes including *PDCD1*, *LAG3*, and *TIGIT* in the overall CD8+ T cell population (Fig. 3H). This suggests that HBV infection promotes CD8+ T cell exhaustion, potentially contributing to immune dysfunction in the HBV-infected environment.

To further explore the clinical impact of T cell subtype distribution, we conducted a survival analysis using TCGA LIHC data, focusing on T cell infiltration levels defined by our scRNA-seq data. Using CIBERSORTx, we found that increased infiltration of CD8-c7-PDCD1 subtype, indicative of exhaustion, was correlated with poorer prognosis (p = 0.014; Fig. 3I). These data were consistent with the observation that the PDCD1+ CD8 T cells displayed higher levels of exhaustion marker expression (Fig. 3H), indicating that inhibiting PDCD1+ CD8 T cells may have positive effect on the overall survival of patients. Since transcriptional regulation is a key determinant in T cell differentiation and function, we used pySCENIC to identify the most active transcription factors (TFs) across T cell subtypes. Especially, we found that for the CD4-c3-IL7R subtype, MXI1 and EOMES showed the highest regulon activity, while for CD8-c7-PDCD1 subtype, RELA, RUNX3, POLR2A, and MAFB showed the highest regulon activity (Fig.3J). Collectively, these data suggest that those important TF regulators may perform regulatory roles on the fate determination or differentiation of the T cells shaped by the HBV-infection status.

In summary, our analysis of T cell heterogeneity in HCC revealed distinct CD4+ and CD8+ T cell subtypes with or without HBV-infection condition and uncovered that certain T cell subtype associated with patient outcomes, such as CD8-c7-PDCD1, linked to poorer survival. These findings, combined with transcription factor regulon analysis, enhance our understanding of T cell diversity in HBV-infected versus non-infected HCC. This knowledge highlights the importance of targeting specific T cell subtypes and regulatory pathways to improve therapeutic strategies for HCC. Moreover, DEG analysis revealed that one gene, SLC35F1 was commonly upregulated in CD4 and CD8 T cells upon HBV-infection. Further research is needed to investigate the role played by SLC35F1 in HBV-infected HCC.

### Characterization of macrophages and dendritic cells in HBV-Infected and Non-HBV-Infected HCC

Macrophages and dendritic cells (DCs) are integral components of the tumor immune microenvironment, playing the key roles in immune regulation and interactions with other immune cells(26). To investigate the heterogeneity of these myeloid populations in HCC, we analyzed scRNA-seq data from HBV-infected and non-HBV-infected samples.

Macrophages are essential for regulating inflammation, tumor immunity, and tissue remodeling, and their phenotype and function may be altered by chronic HBV infection. By unsupervised clustering analysis, we identified five distinct macrophage subtypes (Fig. 4A). The distribution of these subtypes varied between HBV-infected and non-HBV-infected samples, suggesting that HBV infection reshapes the macrophage landscape in HCC (Fig. 4B and Fig.S3D). To better understand the functional roles of these subtypes, we conducted marker gene analysis, which revealed distinct immune functions across the macrophage populations (Fig. 4C). The data showed that the proportion of IFITM3+ macrophages was evidently increased and SPP1+ macrophages was decreased upon HBV-infection, respectively (Fig.4B), which suggest that these two subtypes of macrophages might have some associations with HBV-infection status. To further compare the difference of each macrophage’s subtypes, we performed DEG analysis between HBV-positive and negative groups. Interesting, the results showed that SLC35F1 gene was only significantly upregulated in Macro-c4-NLRP3 subtypes (Fig.4D), which is different from the DEG results in T cells analyzed above, suggesting SLC35F1 gene might be an important mediator or functional gene responsive to HBV infection in this subtype.

**Fig. 4.**
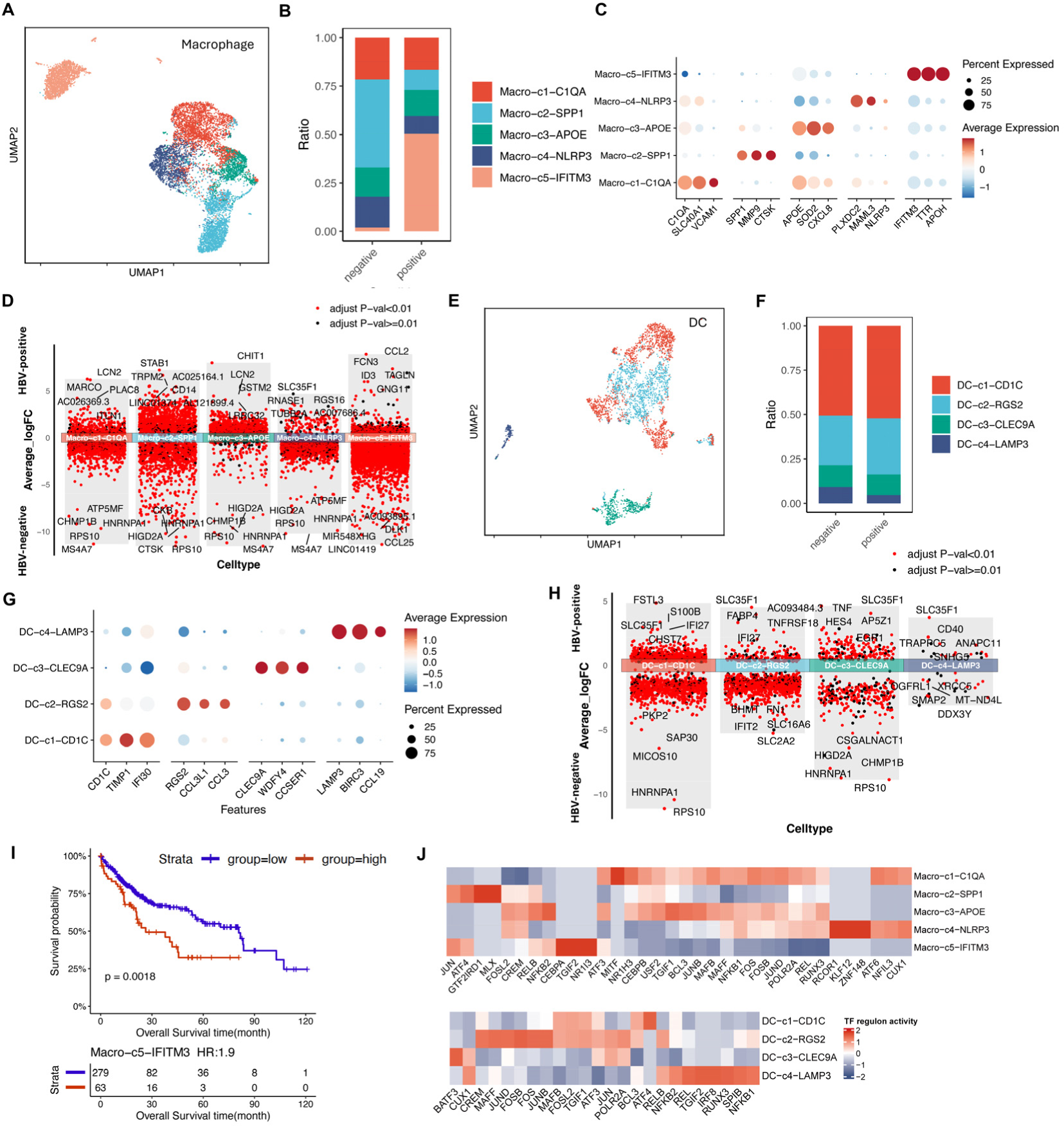
Analysis of macrophage and dendritic cell heterogeneity. (A) UMAP plot of macrophages with subtypes distinguished by color. (B) Stacked bar plots summarizing the proportions of macrophage subtypes in HBV-negative and HBV-positive samples. (C) Dot plot showing the percentages of cells expressing marker genes alongside their average expression levels across macrophage subtypes. (D) Volcano plot highlighting DEGs between HBV-negative and HBV-positive macrophages. (E) UMAP plot of dendritic cells (DC), with subtypes color-coded. (F) Stacked bar plots summarizing the proportions of DC subtypes in HBV-negative and HBV-positive samples. (G) Dot plot showing the percentages of cells expressing marker genes alongside their average expression levels across DC subtypes. (H) Volcano plot of DEGs comparing HBV-negative and HBV-positive DCs. (I) Kaplan–Meier curve illustrating LIHC patient survival based on Macro-c5-IFITM3 infiltration levels. (J) Heatmap showing normalized TF regulon activity in macrophage and DC subtypes as predicted by pySCENIC.

DCs are key antigen-presenting cells that are vital for the initiation and regulation of T cell responses. We classified DCs into four subtypes, representing various maturation stages and specialized functions (Fig. 4E). Differences in the distribution of DC subtypes between HBV-infected and non-HBV-infected samples suggest that HBV infection may impair DC-mediated antigen presentation and subsequent immune activation (Fig. 4F and Fig.S3E). We found that upon HBV-infection, the proportion of RGS2+ DC was increased and LAMP3+ DC was decreased upon HBV-infection, respectively (Fig.4F). The functionality of these DC subtypes was further evaluated by analyzing the expression of key activation and co-stimulatory markers, which demonstrated varying capacities to stimulate anti-tumor immune responses (Fig. 4G). Similar, DEGs analysis was performed to compare the difference of DC subtypes with or with HBV infection. The results highlighted the upregulated gene, SLCF351, upon HBV-infection (Fig.4H).

To assess the clinical relevance of macrophage and DC heterogeneity, we performed a survival analysis using data from the TCGA LIHC dataset. We observed that increased infiltration of the macrophage subtype Macro-c5-IFITM3 correlated with poorer overall survival (p = 0.0018; Fig. 4I). These results suggest that an imbalance favoring immune-suppressive macrophages may negatively affect patient outcomes in HCC. Especially, the evidently high proportion of IFITM3+ macrophage upon HBV-infection may account for the lower survival rate of HCC.

We further explored transcriptional regulation in these myeloid subtypes using pySCENIC to identify active transcription factors (TFs). In the macrophage Macro-c5-IFITM3 subtype, CEBPA, TGIF2, and NR1I3 exhibited high regulon activity, indicating their role in driving specific functional states (Fig. 4J). In DC-c4-LAMP3, TFs such as CUX1, REL, TGIF2, IRF8, RUNX3, SPIB, and NFKB1 were identified as key regulators guiding DC differentiation and function (Fig. 4J).

In summary, our analysis reveals distinct macrophage and DC subtypes in HCC that are associated with patient survival, with subtypes Macro-c5-IFITM3 linked to poorer outcomes. Transcription factor analysis highlighted key regulators that drive these immune cell states, providing a more comprehension of the immune landscape in HBV-infected and non-HBV-infected HCC and identifying potential targets SLC35F1 and IFITM3 for improving immune response in HCC.

### HBV infection shapes the cell-cell communication of HCC

To understand the impact of HBV infection on intercellular communication within HCC samples, the differential cell-cell communication analyses were performed. We identified 22,656 significant ligand-receptor (L-R) interactions in non-HBV-infected HCC and 20,049 in HBV-infected HCC (Fig. 5A, left panel). The total strengths of these interactions were 586.425 and 599.8, respectively (Fig. 5A, right panel). Although the overall number of L-R interactions in tumor liver tissues decreased due to HBV infection, the infection exhibited distinct effects on various cell-cell communication patterns. For instance, interactions from endothelial cells to fibroblasts and Macro-C3-APOE were enhanced in HBV-infected tissues, while the incoming signaling interactions of NK cells decreased (Fig.S4). Additionally, we observed that HBV infection increased incoming signaling for monocytes, dendritic cells (DC), and CD8 T cells (Fig. 5B). Collectively, these findings underscore the substantial impact of HBV infection on intercellular communication within the tumor microenvironment.

**Fig. 5.**
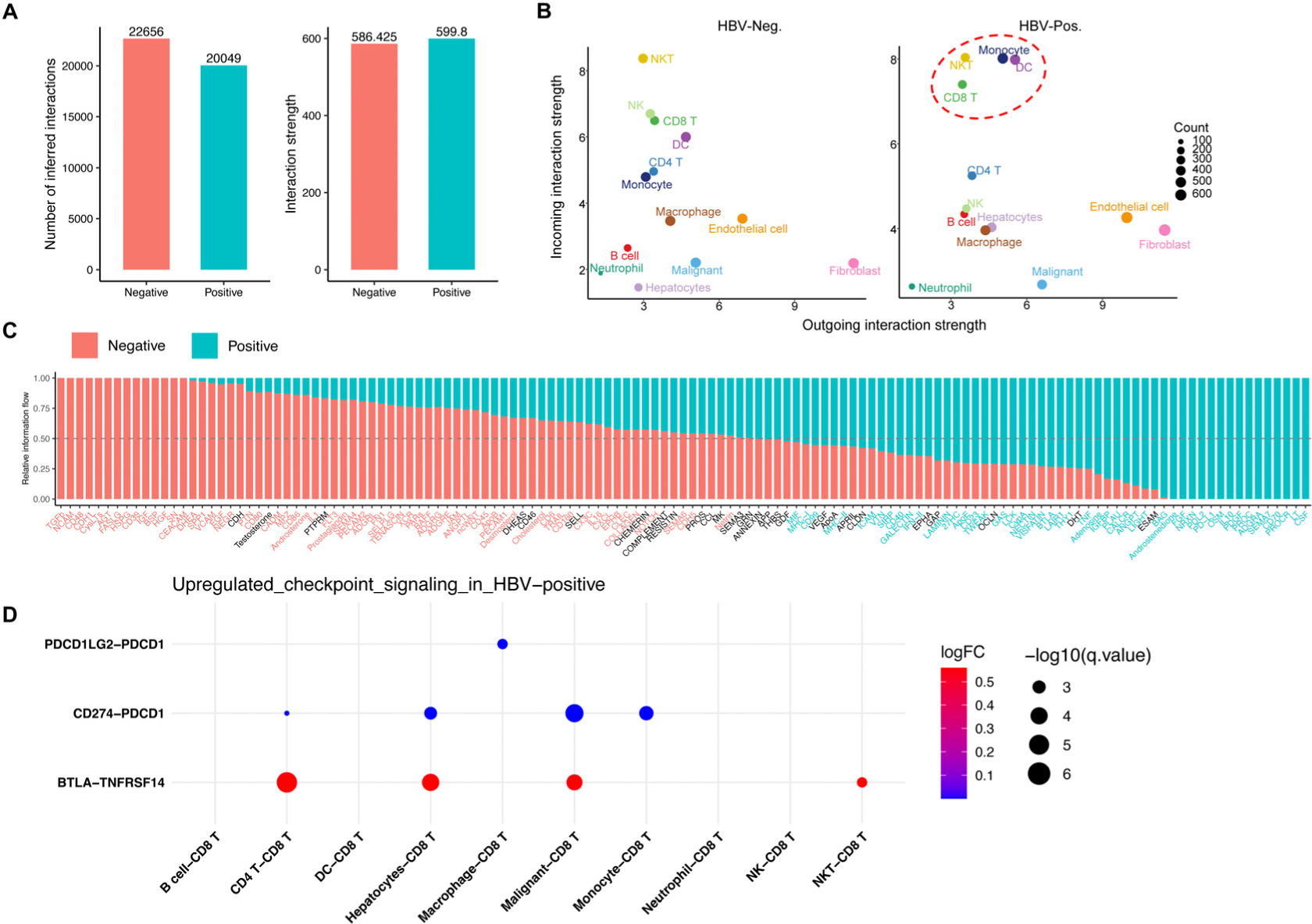
The impact of HBV infection on intercellular communication within HCC. (A) Bar plots comparing the number of cellular interactions (left panel) and their interaction strength (right panel) between HBV-infected (HBV-Pos.) and non-HBV-infected (HBV-Neg.) HCC samples. (B) Dot plots illustrating the strength of outgoing and incoming cellular communications for each cell type in HBV-Pos. (right) and HBV-Neg. (left) samples. Each cell type is represented by a specific color, and the size of the dot reflects the number of cellular interactions. (C) Stacked bar plot depicting the information flow between HBV-negative and HBV-positive conditions. (D) Dot plot highlighting upregulated checkpoint signaling pathways in HBV-positive samples.

We next aimed to explore how HBV infection influences incoming and outgoing signaling by calculating the distance between these pathways based on functional similarity. We identified ESAM, SEMA3, and ANGPT as the top three pathways affected by HBV (Fig. S5). Besides, from the perspective of information flow, the results revealed four types of information flow, HBV-negative specific flow, HBV-negative biased flow, HBV-positive biased flow and HBV-positive specific flow (Fig. 5C), which provided a comprehensive overview of communication signaling flow.

Our analysis previously demonstrated that HBV infection enhances CD8+ T cell exhaustion, as evidenced by the significantly increased expression of exhaustion markers *LAG3*, *PDCD1* and *TIGIT* (Fig. 3H). However, the specific cell types or ligand-receptor signaling pairs mediating this exhaustion process remained unclear. To address this, we investigated the interactions between CD8+ T cells and other immune cells, as well as malignant tumor cells, with a focus on outgoing checkpoint signaling targeting CD8+ T cells. The analysis revealed that three checkpoint signaling pathways—*PDCD1LG2-PDCD1*, *CD274-PDCD1*, and *BTLA-TNFRSF14*—were significantly upregulated under HBV-infected conditions compared to HBV-non-infected conditions (Fig. 5D). Furthermore, the data showed that while CD4+ T cells, hepatocytes, and malignant tumor cells primarily contribute *BTLA* and *CD274* ligands, macrophages are the exclusive source of the suppressive *PDCD1LG2* ligand (Fig. 5D). Collectively, our cell-cell communication analysis suggests that HBV infection contributes to CD8+ T cell exhaustion by increasing suppressive interactions with other cell types in the tumor microenvironment altered by HBV.

## Discussion

The onset and progression of tumors, including hepatocellular carcinoma, are significantly influenced by the immune microenvironment(27, 28). Recently, single-cell sequencing technology has been used to reveal the immune microenvironment characteristics of various tumors, providing new strategies for immunotherapy(16-20, 25, 29). In the context of HCC, several single-cell omics studies explore the immune profile within tumor, revealing new T-cell subsets and demonstrating the immunosuppressive environment and tumor heterogeneity(18, 19, 25, 30). It is noteworthy that HCC patients often have HBV infections, a risk factor that inevitably affects the immune system(6). Theoretically, HBV infection alters the immune microenvironment, which will impact tumorigenesis and progression. However, the previous studies have not thoroughly addressed this important issue. In this study, we performed a comparative analysis of the immune microenvironments in HCC patients with and without HBV infection. We found that HBV infection significantly alters the tumor immune microenvironment, placing it in a more suppressed and exhausted state. This discovery provides valuable insights into the immunological mechanisms by which HBV infection contributes to HCC development, paving the way for immunotherapy strategies.

We found a marked increase in the proportion of MAIT cells, characterized by the expression of SLC4A10 and CD8, among the infiltrating T cells in HBV-positive HCC versus HBV-negative HCC. Unlike conventional αβ+ CD8+ T cells that detect peptide antigens via MHC class I, MAIT cells recognize metabolites derived from microbes, which are presented by the MHC class I-related protein MR1(31, 32). While previous reports have showed the potential antitumor activity of MAIT cells, demonstrating their cytolytic activity against tumor cells in vitro under certain conditions(33-35), recent in vivo findings suggest that MAIT cells could actually facilitate the initiation, growth, and metastasis of tumors(36, 37). This tumor-promoting effect is believed to result from MAIT cells inhibiting the functions of T cells and/or NK cells. Therefore, we hypothesize that the rise in MAIT cell proportion triggered by HBV infection may contribute to the progression of HCC. Consequently, targeting MAIT cells in HBV-associated HCC could potentially represent a novel immunotherapeutic strategy.

Interestingly, we identified a gene, SLC35F1, which is significantly upregulated in various immune cells within HBV-positive HCC. Research on SLC35F1 is limited, and its specific function remains unclear. However, it belongs to the SLC35 family, which mediates the transport of nucleotide sugars across biological membranes(21, 22), suggesting that SLC35F1 may play a potential role in the transport and metabolism of nucleosides. In the complex tumor microenvironment, nucleotide metabolism profoundly influences various immune cells, modulating their functions and shaping the cancer immunity landscape(38, 39). Consequently, SLC35F1 may be involved in regulating T cell function through nucleotide metabolism. Although the specific role of SLC35F1 requires further investigation, it is possible that the influence of HBV infection on the tumor immune microenvironment is mediated through SLC35F1. Therefore, SLC35F1 could be a potential target for modulating the immune microenvironment in HBV-positive HCC.

There are several limitations to this study. First, the sample size is somewhat small, comprising only three cases each of HBV-positive and HBV-negative patients. Additionally, all hepatocellular carcinoma patients included in this study were at TNM stage I, which may limit the generalization of our findings and conclusions to patients with more advanced stages of liver cancer. Second, our research is based on single-cell RNA sequencing data; therefore, future studies will be necessary to validate these findings through wet lab experiments using patient samples to enhance the robustness of the conclusions.

## Acknowledgments

Not applicable.

## Funding

This work was supported by Grant 2023YFA0915400 (to H.L.) from the National Key R&D Program of China, Grants 32371450 (to X.S.) and 82303446 (to E.C.) from the National Natural Science Foundation of China, Grants 2024A1515011562 (to K.L.), 202381515040008 (to H.L.) and 2023A1515220200 (to E.C.) from the Natural Science Foundation of Guangdong Province, Grant A2024351 (to E.C.) from Medical Scientific Research Foundation of Guangdong Province, Grants JCYJ20210324120200001 (to H.L.), JCYJ20210324101805014 (to K.L.) and JCYJ20200109114608075 (to X.S.) from Shenzhen Science and Technology Program, Grant KYQD2023303 (to E.C.) from Shenzhen High-level Hospital Construction Fund, and Peking University Shenzhen Hospital Scientific Research Fund.

## Availability of data and materials

All scRNA-seq data are available in the Gene Expression Omnibus database with the accession code GSE282701. Additional data supporting this study’s findings can be found within the article or obtained from the corresponding authors upon request.

## Author contributions

H.L., X.S., and D.Y. conceived the project and supervised the study. E.C. collected specimens and clinical information. K.L., and J.L. prepared samples for scRNA-seq. K.L., J.L., and Y.L. analyzed scRNA-seq data. B.C., W.S., Z.Z., W.Z., and H.T. provided special technical support and discussed the data. K.L., X.S., D.Y. and H.L. wrote the manuscript.

## Ethics approval and consent to participate

All the involved patients have signed the consent informs, and this study was approved by the Ethics Committee of Peking University Shenzhen Hospital (Approval No. 2022-164).

## Competing interests

The authors declare no competing interests.

## Supplementary Figure Legends

**Fig.S1 Sample information and clustering annotation in primary HCC and non-tumor liver tissues.**

(A) Sample information sheet of HCC or HCC-adjacent tissues from 6 HCC patients. (B) Violin plot showing the detected genes across the samples after quality control. (C) The UMAP plots showing the identification of 11 clusters with unsupervised clustering method. (D) Dotplot showing the marker genes expression of 11 clusters.

**Fig. S2 Further clustering of HBV-infected vs. non-HBV-infected HCC**

(A) Stacked bar plots summarizing the proportions of major cell types in HBV-negative and HBV-positive samples. (B) Dot plot displaying the expression of marker genes across 13 annotated clusters. (C) Bar plot showing upregulated and downregulated GO terms associated with HBV infection. (D) Dot plot highlighting the upregulated and downregulated KEGG pathways in HBV-infected samples. (E) GSEA plot illustrating altered KEGG pathways between HBV-positive and HBV-negative samples.

**Fig. S3 Proportions of cell subtypes across individual patients** (A) Bar plots illustrating the distribution of CD4+ T cell subtypes across each sample. (B) Bar plots depicting the distribution of CD8+ T cell subtypes across each sample. (C) Dot plot displaying the percentages of cells expressing marker genes alongside their average expression levels across CD8+ T cell subtypes. (D) Bar plots showing the proportions of macrophage subtypes across each sample. (E) Bar plots presenting the proportions of dendritic cell (DC) subtypes across each sample.

**Fig.S4 The effect of HBV infection on intercellular communication within tumor liver tissues across all identified cell subtypes.**

Increased intercellular communication is represented in red, while decreased communication is shown in blue.

**Fig.S5 Identifying the effect of HBV infection on signaling networks within tumor liver tissues.**

Bar plot showing the pathway distance in HBV Pos. versus HBV Neg. Only pathway distances of more than 1 were shown.

## Notes

### Competing Interest Statement

The authors have declared no competing interest.

