## Supplementary figures and images for "Single-cell analysis reveals distinct immune characteristics of hepatocellular carcinoma in HBV-positive versus HBV-negative cases"

### Figure S1

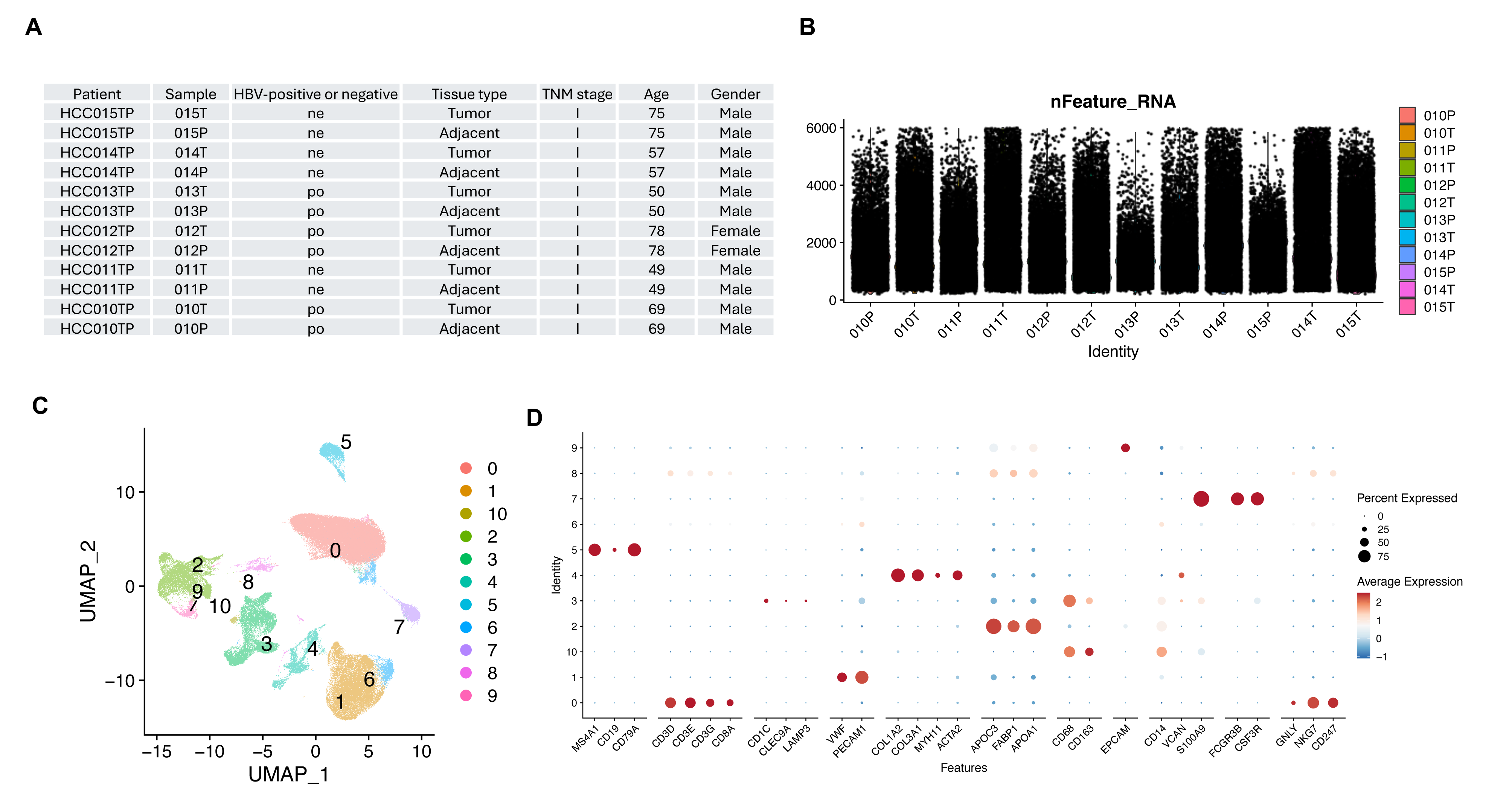

### Figure S2

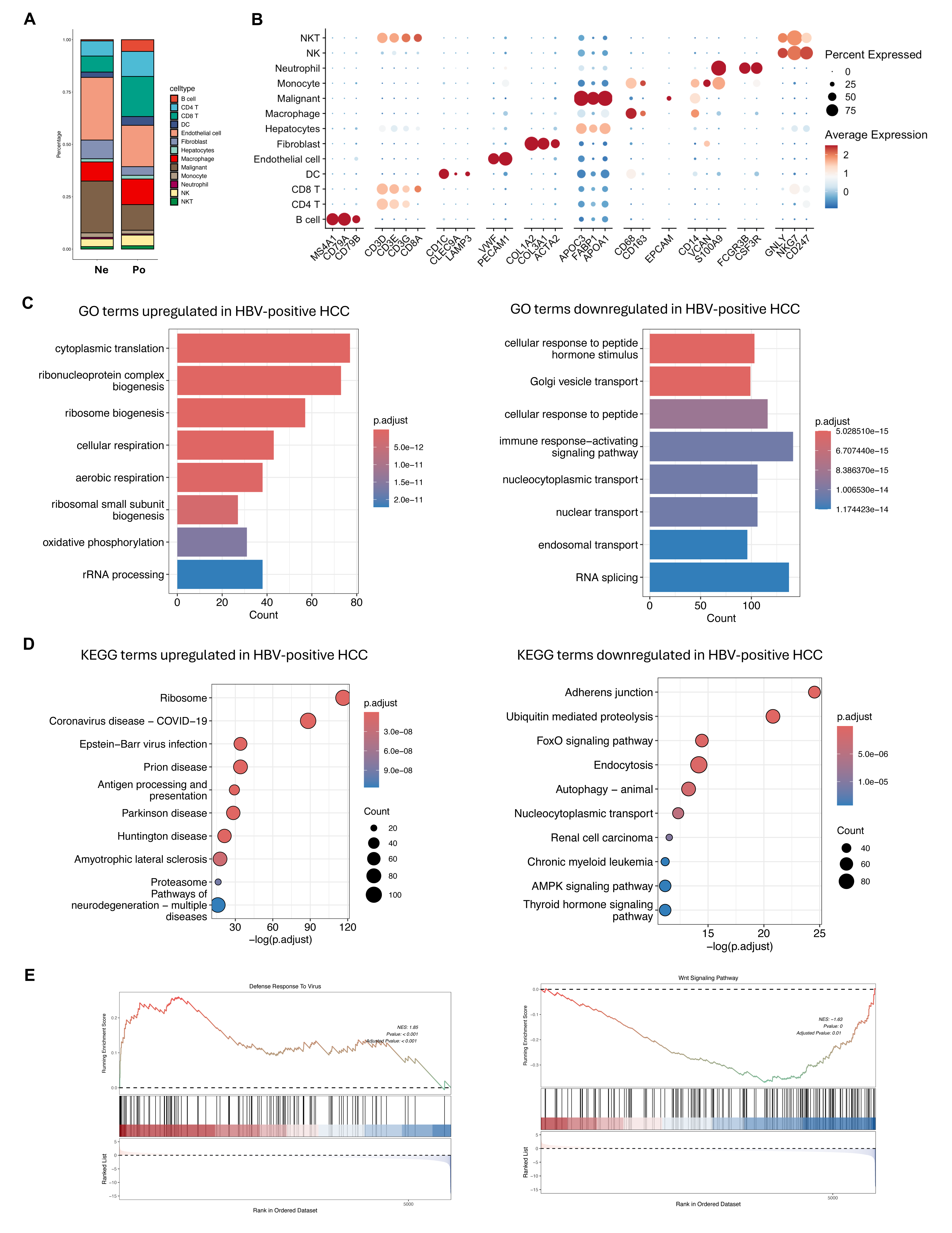

### Figure S3

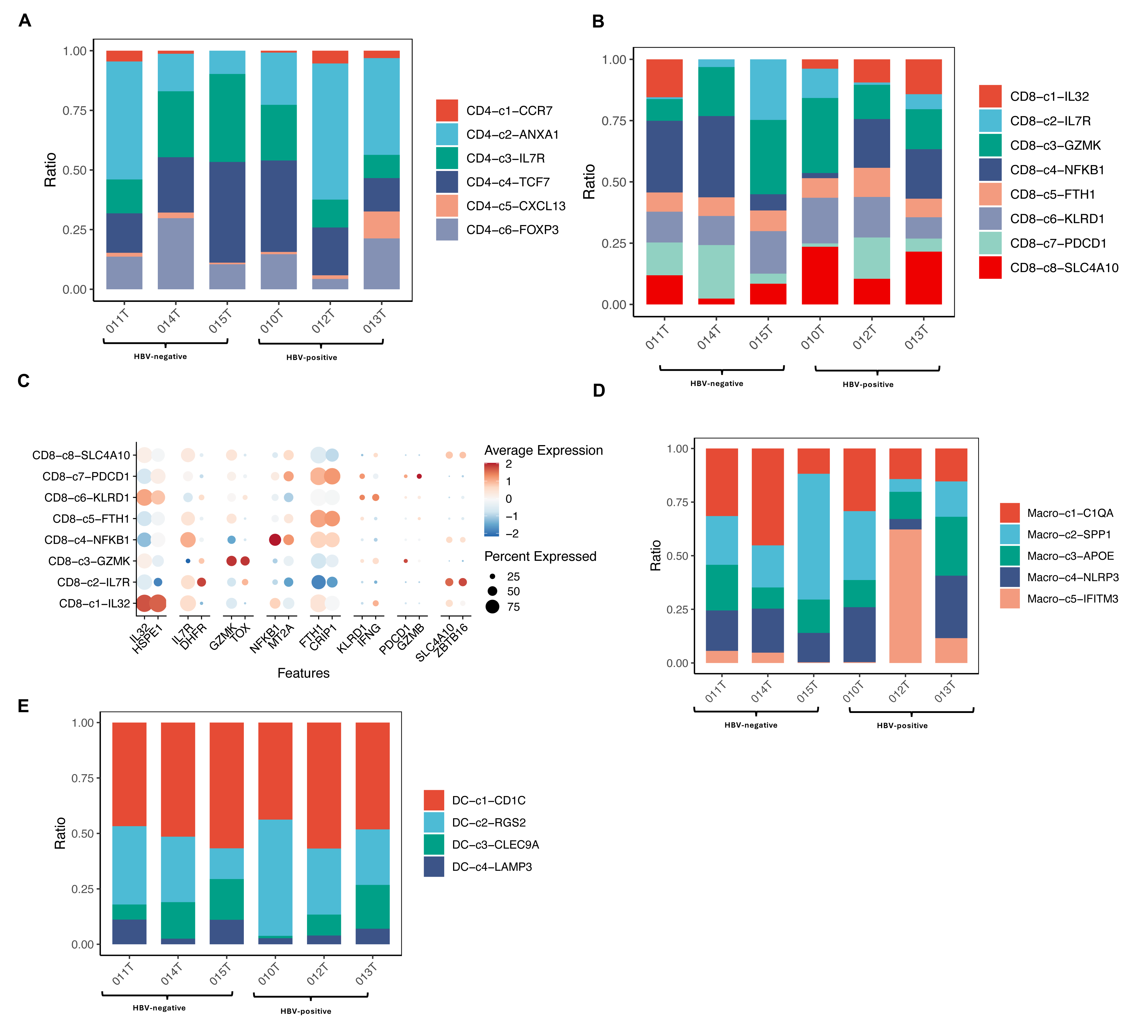

### Figure S4

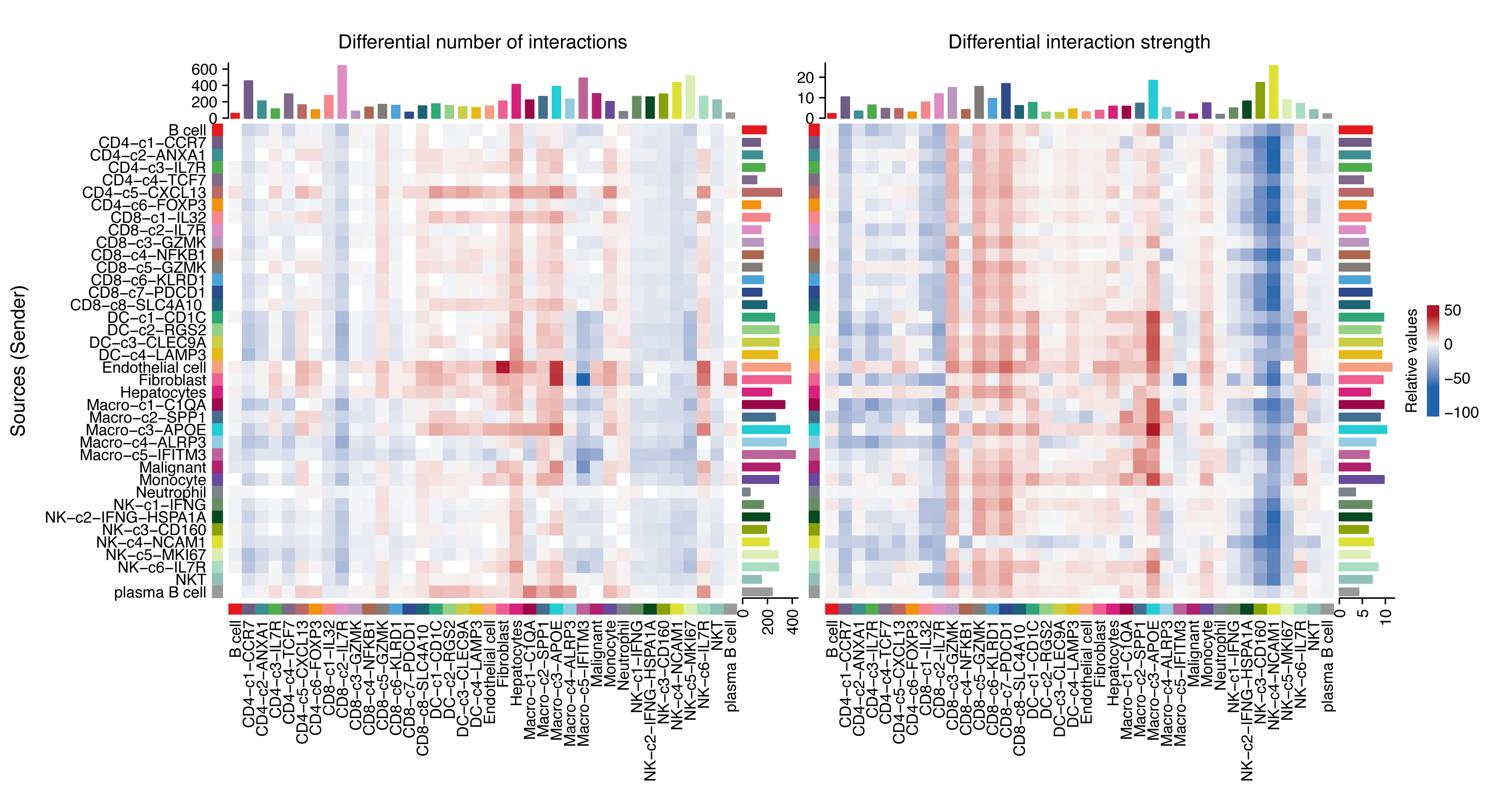

### Figure S5

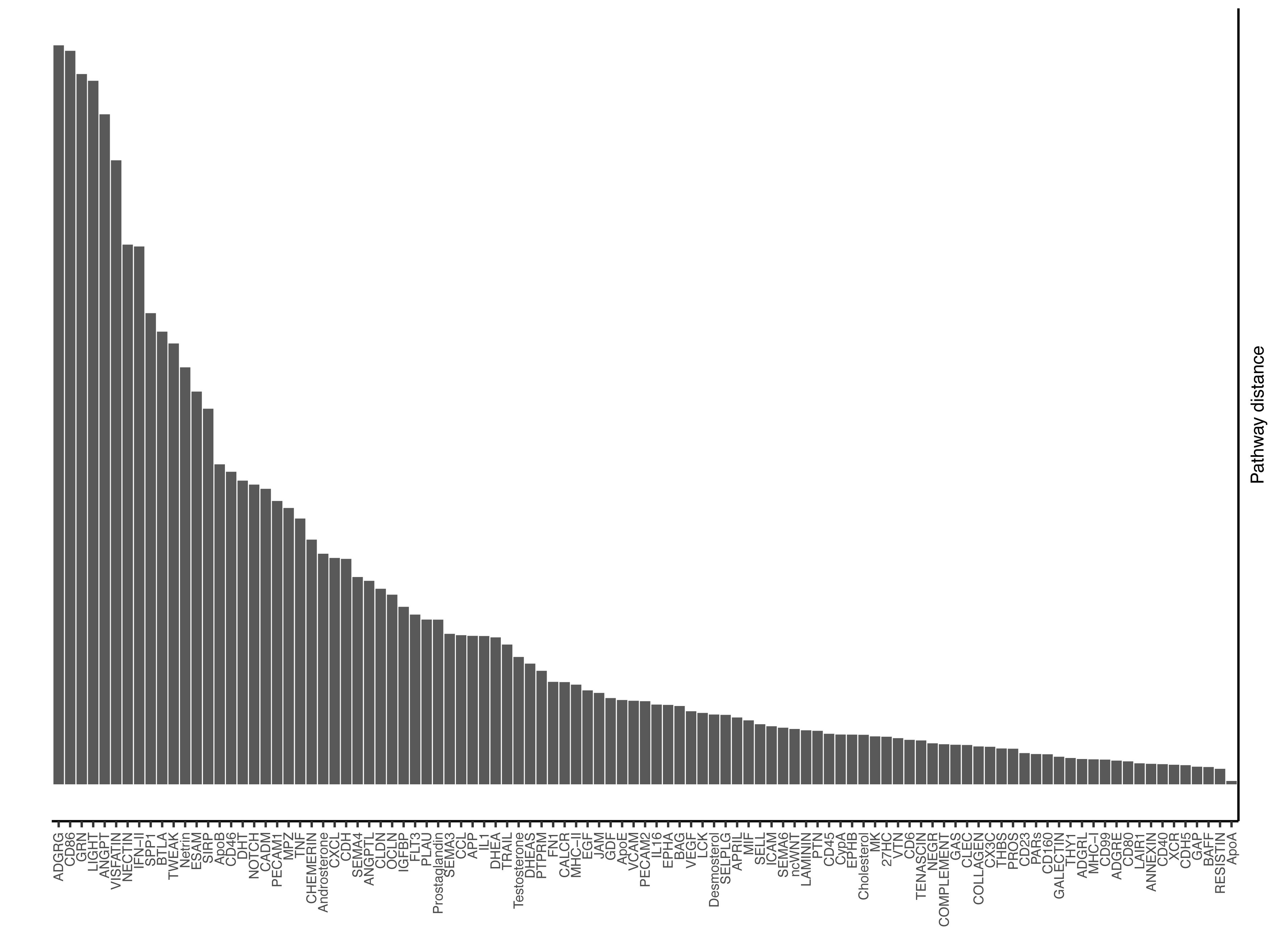
